# Establishment of Bovine Trophoblast Stem Cells

**DOI:** 10.1101/2022.12.21.521294

**Authors:** Yinjuan Wang, Leqian Yu, Jie Li, Linkai Zhu, Hao Ming, Carlos Pinzon Arteaga, Hai-Xi Sun, Jun Wu, Zongliang Jiang

**Author notes:** To whom correspondence will be addressed.

## Abstract

Here we report that a chemical cocktail (LCDM: h<u>L</u>IF, <u>C</u>HIR99021, <u>D</u>iM and <u>M</u>iH) previously used for extended potential pluripotent stem cells enables the de novo derivation and long-term culture of bovine trophoblast stem cells (TSCs). Bovine TSCs exhibit transcriptomic and epigenetic features characteristic of trophectoderm cells from bovine embryos and retain developmental potency to differentiate into mature trophoblast cells.

## Introduction

Trophoblasts are specialized cells in the placenta that mediate maternal-fetal crosstalk and are originated from the trophectoderm (TE) of the blastocyst. Pregnancy establishment in cattle requires TE elongation, a unique process in ruminants prior to apposition, attachment, and implantation [1]. Although proper placental development and function are pivotal for gestational success and the failure of which often results in a range of adverse pregnancy outcomes [2-4], there lacks *in vitro* models to study ruminant trophoblast development. Trophoblast stem cells (TSCs) have been established from several rodent and primate species including mice [5], humans [6], and nonhuman primates [7]. Despite several attempts [8-10], however, bona fide bovine TSCs that withstand the rigor of long-term culture have yet to be derived.

## Results

TE cells of bovine blastocysts retain the plasticity to generate ICM cells, and vice versa [11, 12], which prompted us to test de novo derivation of bovine TSCs with different combinations of basal media, growth factors and chemicals that were previously used for culturing pluripotent stem cells (PSCs) (**Extended Data Table. 1**). We identified four conditions that could support bovine blastocyst outgrowth for several passages on mouse embryonic fibroblasts (MEF) feeder cells (**Fig. 1a, b and Extended Data Fig. 1a**). Interestingly, an extended pluripotent stem cell (EPSC) culture condition, LCDM (h<u>L</u>IF, <u>C</u>HIR99021, <u>D</u>iM and <u>M</u>iH) [13], was most effective to support long term passage (more than 70 passages at the time of writing) of bovine TSC-like cells (bTSC-LCs) from blastocyst outgrowth. Removing each of hLIF, CHIR99021, DiM and MiH failed to maintain the morphology and self-renew of bTSC-LCs (data not shown). bTSC-LCs could also be maintained feeder-free on Matrigel in the presence of MEF-conditioned LCDM medium (**Fig. 1b**). Further characterization revealed that: 1) bTSC-LCs maintained stable colony morphology and a normal diploid number of chromosomes (60) after long-term *in vitro* culture (**Fig. 1b and Extended Data Fig. 1b**). 2) bTSC-LCs highly expressed TE-related transcription factors (TFs) (*CDX2, SFN, ELF3, GATA3, ASCL2, GATA2* and *ETS2*) (**Extended Data Fig. 1c**). 3) Similar to TE cells in bovine blastocysts, at the protein level, bTSC-LCs expressed CDX2, GATA3 and KRT8 but not SOX2 (**Fig. 1c and Extended Data Fig. 1d**). 4) The majority of bTSC-LCs were found GATA3+ (**Fig. 1d**). Collectively, these results demonstrate that the LCDM medium supports de novo derivation and long-term self-renewal of bTSC-LCs.

**Fig. 1.**
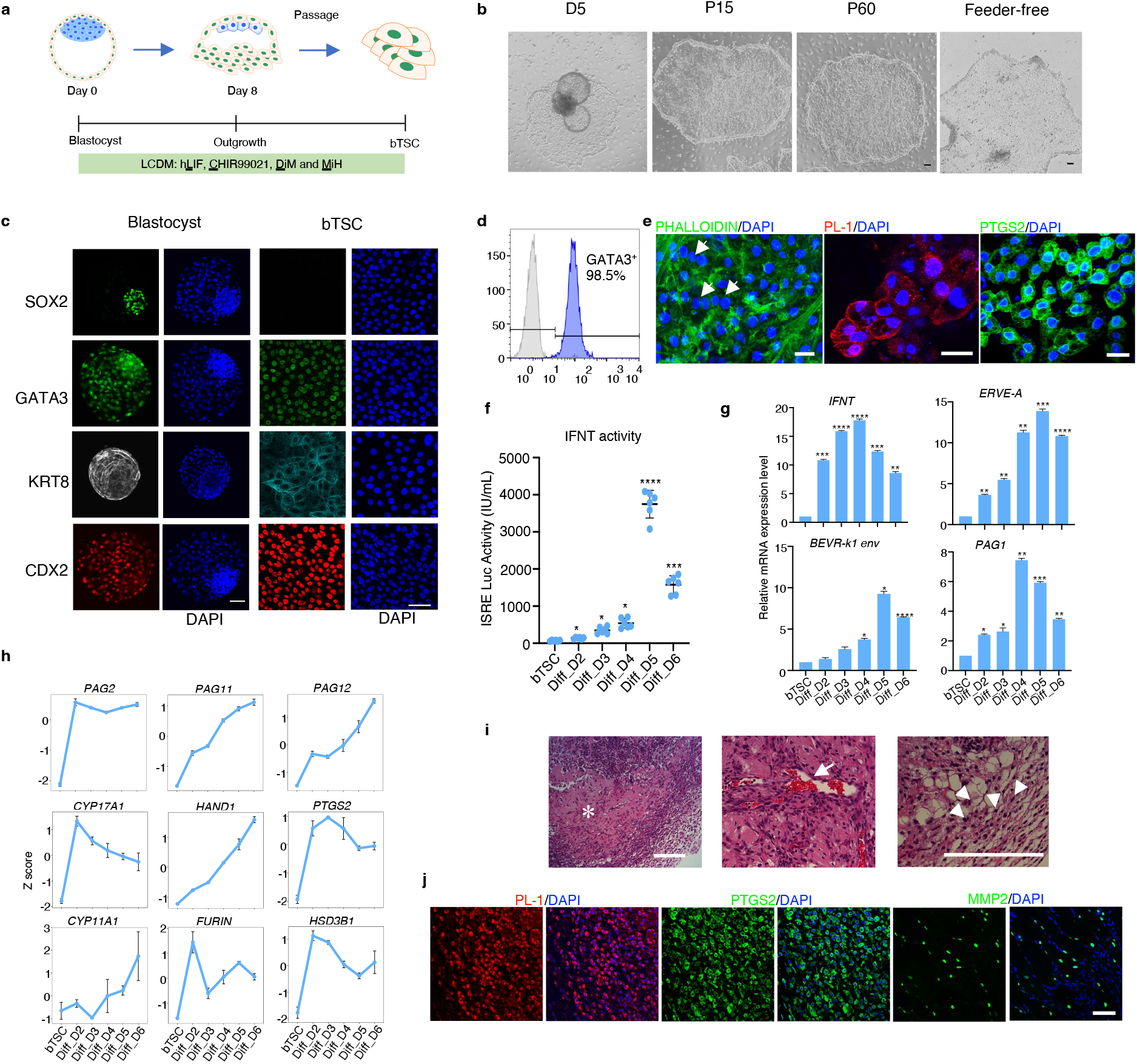
Derivation and characterization of bovine TSC. **a**, Illustration of the derivation of bovine TSC from blastocyst. **b**, Bright field images of the outgrowths of blastocysts and typical morphologies of bovine TSC on feeder or feeder-free. D5: outgrowth after 5 days culture; P15: passage 15; P60: passage 60. Scale bar: 100 μm. **c**, Immunostaining for epiblast marker SOX2, and trophoblast marker (GATA3, KRT8 and CDX2) in bovine Day 7 IVF blastocysts and bTSC. (Scale bar: 50 μm). **d**, Flow cytometry quantification of GATA3 positive cell population in bTSC. **e**, Immunostaining for bovine mature trophoblast markers (PL-1 and PTGS2) in differentiated-bTSC (P55). Scale bar: 25 μm. **f**, IFNT activity secreted by bTSC and trophoblast cells differentiated from bTSC from Day 2 to Day 6 (n=6). IFNT: interferon tau. **g**, Expression levels of *IFNT, ERVE-A, BEVR-k1 env* and *PAG1* during *in vitro* differentiation. **h**, Expression dynamics of mature trophoblast cell marker genes (*PAG1, PAG11, PAG12, CYP17A1, HAND1, PTGS2, CYP11A1, FURIN* and *HSD3B1)* during bTSC in vitro differentiation. **i**, H&E staining analysis of bovine TSC-derived lesion. Asterisk: necrotic area; Arrow: blood-filled *lacunae;* Arrowhead: binucleate cells. Scale bar: 200 μm. **j**, Immunostaining for mature trophoblast markers (PL-1, PTGS2) and trophoblast-endometrial regulator (MMP2) in TS-derived lesion. Scale bar: 75 μm.

We next assessed the differentiation potential of bTSC-LCs *in vitro*. We found a condition containing forskolin, Y27632 and 4% knockout serum replacement (KSR) enabled the differentiation of bTSC-LCs into bi-nucleated cells, which expressed trophoblast markers PTGS2 and placental lactogen 1 (PL-1) (**Fig. 1e and Extended Data Fig. 1e, f, g**). In ruminants, interferon tau (IFNT) produced by mature trophoblast cells is known as a signal for maternal recognition of pregnancy [14]. By using a luciferase-based IFN stimulatory response element (ISRE) assay [15], we found IFNT production significantly increased upon differentiation of bTSC-LCs and peaked around day 5 (**Fig. 1f**). qRT-PCR analysis further showed that the expression of *IFNT* and mature trophoblast markers (*BEVR*-*k1 env, bEPVE*-*A* [16] and pregnancy associated glycoproteins 1 (*PAG1*) [17]) were significantly upregulated following bTSC-LCs differentiation (**Fig. 1g**). We performed RNA-sequencing (RNA-seq) across six timepoints during bTSC-LCs differentiation and found bTSC-LCs transitioned through an intermediate stage on day 2 before further differentiation into more mature trophoblast cells between days 3-6 (**Extended Data Fig. 2a**). As expected, RNA-seq analysis showed that PAG family genes (*PAG2, PAG11* and *PAG12*) and well-known bovine placental markers (*CYP11A1, CYP17A1, FURIN, HAND1, PTGS2* and *HSD3B1*) [17-19] were upregulated during differentiation (**Fig. 1h**). Differentiated trophoblasts (day 4) had an up-regulation of genes with their gene ontology (GO) terms related to morphogenesis, cell migration, and locomotion (**Extended Data Fig. 2b**), suggesting the presence of invasive trophoblast cells. In addition, differentiated trophoblast cells expressed a number of genes involved in signaling pathways such as ECM-receptor interaction, TNF, IL-17, and MAPK signaling (**Extended Data Fig. 2c**), which is consistent with the increase of these signaling activities during implantation and placental development in ruminants and humans [20-22].

We also determined the differentiation potential of bTSC-LCs by subcutaneously injecting them into NOD-SCID mice. By day 9, the injected bTSC-LCs formed ∼0.5 cm lesions (**Extended Data Fig. 2d**). Immunohistological analysis revealed that the central area of the lesions was necrotic, and the lesions contained blood-filled lacunae-like structures (**Fig. 1i**), similar to the lesions formed by mouse [23] and human TSCs [6]. Binucleated cells were identified in the peripheral regions of the lesions and expressed PL-1 and PTGS2, suggesting trophoblast maturation (**Fig. 1i and j**). We also identified MMP2 (a key factor for trophoblast-endometrial epithelia crosstalk and remodeling of endometrial matrices [24]) positive cells at the margins of the lesions (**Fig. 1j**). Together, these data demonstrate that bTSC-LCs can generate mature trophoblast cells *in vitro* and *in vivo*, and hereafter we refer them as bTSCs.

We compared the transcriptomes of bTSCs with those from early placental cell types at two different developmental stages: TE of pre-implantation blastocysts (D7_TE) and day 14 embryos (Day 14_TE) [25], and two types of pluripotent stem cells: bovine expanded potential stem cells (bEPSCs) [26] and bovine primed ESCs (bESCs) [27] (**Fig. 2a**). Principal component analysis placed bTSCs in between day 7_TE and day 14_TE, and away from bESC and bEPSCs (**Fig. 2a**). Additional transcriptomic comparisons of TSCs and ESCs among cattle [27], humans [6, 28] and mice [29, 30] confirmed the lineage identity of bTSCs (**Fig. 2b**). bTSCs highly expressed two pluripotency markers *LIN28A* and *SALL4* (**Fig. 2c**), and trophoblast markers *KRT7, TEAD3, ELF3, CDX2*, and *TFAP2A*, which is in contrast with TE cells of early embryos (**Fig. 2c**). In addition, bTSC transcriptomes were enriched with GO terms including intracellular transport and metabolic process (**Extended Data Fig. 3a**) when compared to D7_TE and D14_TE, and hippo signaling pathway, lysosome and tight junction when compared to bESCs and bEPSCs (**Extended Data Fig. 3b**). Of note, signaling pathways including focal adhesion and HIF-1 were also uniquely enriched in bTSCs (**Fig. 2d**).

**Fig 2.**
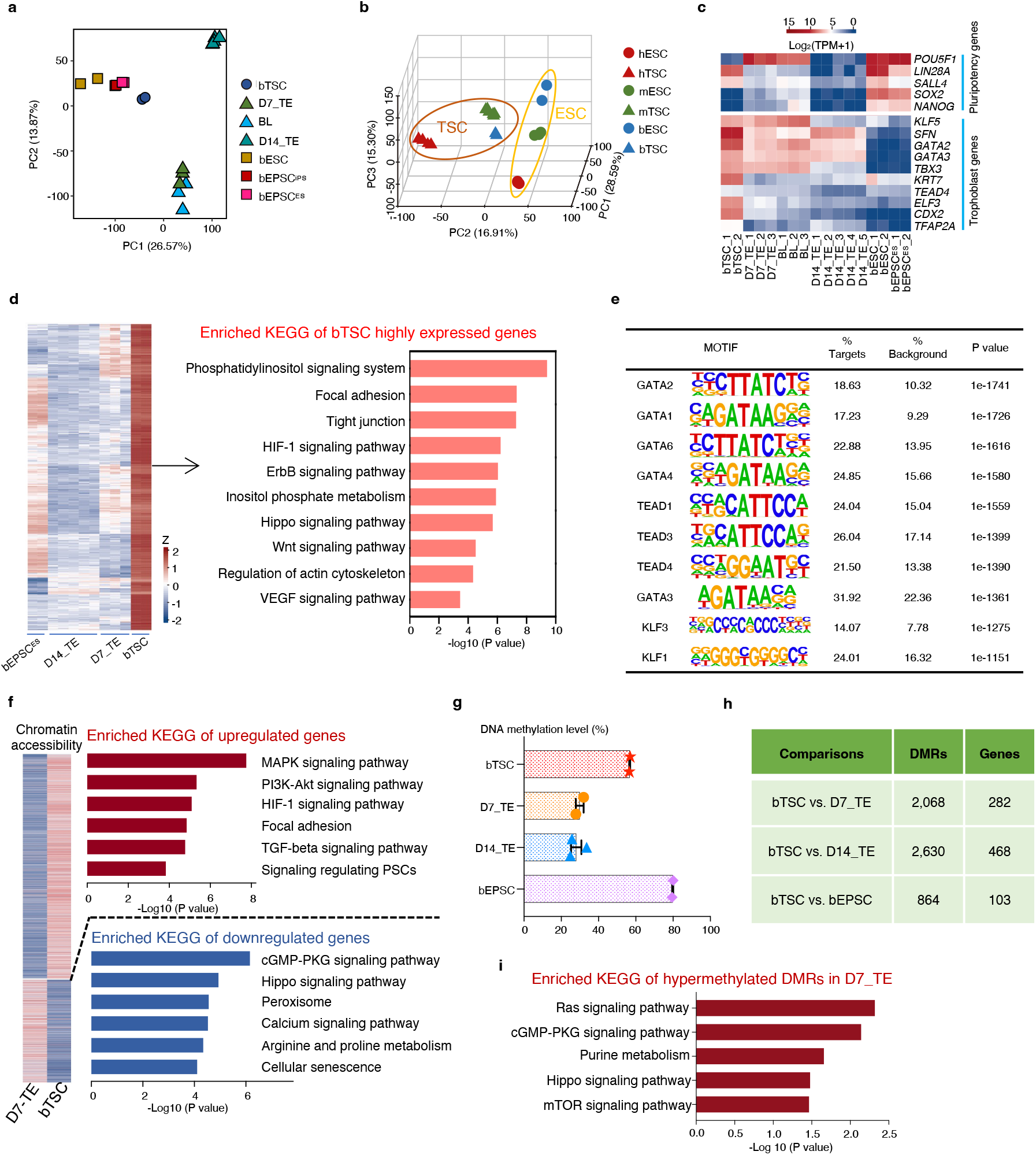
Transcriptomic and epigenomic features of bovine TSC. **a**, PCA analysis of transcriptomes of bTSC, trophectoderm of day 7 IVF blastocyst (D7_TE), and day 14 embryos (D14_TE), day 7 IVF blastocyst (BL), bovine expanded potential stem cells (bEPSCs) and bovine primed ESCs (bESCs). **b**, PCA analysis of transcriptomes of ESC and TSC from three species, human, mouse, and bovine. **c**, Expression pattern of trophoblast and pluripotency marker genes in bTSC, D7_TE, BL, D14_TE, bESC and bEPSC. **d**, Heatmap showing highly specifically expressed genes in bTSC (left panel), and their enriched KEGG pathways (right panel). **e**, Motif enrichment analysis of ATAC-seq peaks from bTSC. **f**, Pathways enriched in genes with more accessible or closed chromatin in bTSC compared to D7_TE. **g**, Average genome-wide DNA methylation levels of bTSC, D7_TE, D14_TE and bEPSC (bTSC, D7_TE and bEPSC: n=2; D14_TE: n=3). **h**, The total number of identified differentially methylated regions (DMRs) and their annotated genes between bTSCs and D7_TE or D14_TE. **i**, Enriched pathways associated with genes annotated from hypomethylated DMRs in bTSC compared to D7_TE.

We also revealed epigenomic features of bTSCs by ATAC-seq and whole genome bisulfite sequencing (WGBS) analyses. We confirmed that trophoblast TFs were among top enriched binding motifs in bTSCs (**Fig. 2e**). Analysis of differential enrichment of ATAC-seq peaks between bTSCs and D7_TE/D14_TE further confirmed the overrepresentation of focal adhesion and HIF-1 signaling pathway in bTSCs (**Fig. 2f, and Extended Data Fig. 3c**). WGBS analysis showed that the overall methylation level of bTSCs (56.75%) were much higher than that of D7_TE (29.90%) and D14_TE (28.03%), but lower than that of bEPSCs [26] (79.80%) (**Fig. 2g**). This is in line with the higher levels of DNA methyltransferases (*DNMT1, DNMT3A* and *DNMT3B)* in bTSCs and bEPSCs (**Extended Data Fig. 3d)**. We were able to identify differentially methylated regions (DMRs) between bTSCs and D7_TE/D14_TE (**Fig. 2h**). Of note, hypomethylated regions in bTSCs included genes that were involved in Ras, cGMP-PKG, calcium signaling and purine metabolism (**Fig. 2i and Extended Data Fig. 3e**). Together, our RNA-seq, ATAC-seq and WGBS analyses provided comprehensive transcriptome and epigenome profiles of bTSCs and shed lights on the molecular features during the earliest steps of placenta development in bovine.

## Conclusion

In this study, we demonstrated that an EPSC culture condition (LCDM) could support de novo derivation of stable bovine TSCs from blastocysts. LCDM-derived bovine TSCs showed the capacity to self-renew long-term in culture while retained the potential to differentiate into mature trophoblast cells. Comprehensive transcriptome and epigenome analyses of bovine TSCs and TEs revealed the molecular features during bovine early placenta development and predicted key regulators of bovine trophoblast differentiation, filling a gap and adding invaluable information of placenta development of an ungulate species. The bovine TSCs established in this study not only serves a model to study the unique placentation process in the ruminants and early pregnancy failure, but also enables the first generation of blastocyst-like structures (blastoids) from a large livestock species (Arteaga et al., manuscript co-submitted).

## Materials and Methods

### Bovine IVF embryo production

The IVF embryos used in this study were produced as previous described [31]. Briefly, bovine cumulus-oocyte complexes (COCs) were aspirated from selected follicles of slaughterhouse ovaries. BO-IVM medium (IVF Bioscience) was used for oocyte in vitro maturation. IVF was performed using cryopreserved semen from a Holstein bull with proven fertility. Embryos were then washed and cultured in BO-IVC medium (IVF Bioscience) at 38.5°C with 6% CO2. Day 7 blastocysts were collected with the zona pellucida removed and were processed for TSC derivation.

### Bovine day 14 elongated embryo production

Day 14 elongated embryo were collected from Holsten cows as previously described [31]. Briefly, superovulation was achieved using five doses of intramuscular injections of FSH beginning five days after insertion of a Controlled Intra-vaginal Drug Release (CIDR) device. Two doses of prostaglandin F2 alpha were given along with the last two FSH treatments, followed by CIDR removal. Standing estrus (Day 0) was seen approximately 48h post-prostaglandin injection. GnRH was then administered at estrus. Each cow was inseminated 12- and 24-hours post-standing heat. Elongated embryos were collected by routine non-surgical uterine flushing on day 14 (D14).

### Derivation and culture of bovine TSCs

Each blastocyst was placed in a separate well of a 12-well plate that was seeded with mitomycin C-treated mouse embryonic fibroblast (MEF) cells. The embryos were cultured in bovine TSC medium containing DMEM: F12 (Gibco) and Neurobasal medium (Gibco) (1:1), 0.5x N2-supplement (Gibco), 0.5x B27-supplement (Gibco), 1x NEAA (Gibco), 1x GlutaMAX (Gibco), 0.1 mM 2-mercaptoethanol (Gibco), 0.1% BSA (MP biomedicals), 10 ng/mL LIF (Peprotech, 300-05), 3 μM CHIR99021 (Sigma, SML1046), 2 μM Dimethinedene maleate (DiM) (Tocris, 1425) and 2 μM Minocycline hydrochloride (MiH) (Santa cruz, sc-203339). All the cells were cultured at 38.5 °C and 5% CO_2_. After 48 hours of plating, the unattached embryos were pressed against to the bottom of the plates with needles under microscope. The culture medium was changed daily. At day 7 or 8, outgrowths were dissociated by Dispase (STEMCELL Technologies) for 5-10 mins at 38.5 °C, followed by twice washes with DMEM/F12. bTSC were passaged mechanically under a microscope. For optimal survival rate, 10 μM Rho-associated protein kinase (ROCK) inhibitor Y-27632 (Tocris, 1254) was added to the culture medium for 24 hours.

Once established, bTSCs were passaged every 6 days at a 1:6 split ratio using Accutase (Gibco, A1110501). Each well of bTSCs was dissociated by 1 mL Accutase for 5 mins at 38.5 °C, the same volume of bTSCs medium was used to dilute Accutase for neutralizing the reaction. bTSCs were cryopreserved by ProFreeze Freezing medium (Lonza, 12-769E) according to the manufacturer’s instructions.

For feeder free condition, bTSCs cultured on feeder cells were passaged to Matrigel (Corning, 354234)-coated plates using MEF-conditioned-bTSC-medium (MEF-bTSC).

### Differentiation of bovine TSCs

Bovine TSCs were grown to 80-90% confluence in the bTSCs medium and dissociated with TrypLE (Gibco, 12605-010) for 15 min at 38.5 °C. Then, bTSCs were seeded in a 6-well plate which was coated with 2.5 μg/mL Col IV (Corning, 354233) at a density of 1 -1.5 × 10^5^ cells per well and cultured in 2 mL differentiation medium containing DMEM: F12 and Neurobasal medium (1:1), with 0.5x N2-supplement, 0.5x B27-supplement, 1x NEAA, 1x GlutaMAX, 0.1 mM 2-mercaptoethanol, 0.1% BSA, 2.5 μM Y27632, 2 μM Forskolin (Sigma, F3917) and 4% KSR (Invitrogen, 10828028). The medium was changed every two days.

### Immunofluorescence analysis

Cells or blastocysts were fixed in 4% paraformaldehyde (PFA) for 20 min at room temperature, and then rinsed in wash buffer (0.1% Triton X-100 and 0.1% polyvinyl pyrrolidone in PBS) for three times. Following fixation, cells were permeabilized with 1% Triton X-100 in PBS for 30 min and then rinsed with wash buffer. Cells were then transferred to blocking buffer (0.1% Triton X-100, 1% BSA and 0.1 M glycine) for 2 hours at room temperature. Subsequently, the cells were incubated with the primary antibodies overnight at 4 °C. The primary antibodies used in this experiment include anti-SOX2 (Biogenex, an833), anti-CDX2 (Biogenex, MU392A; 1:200), anti-GATA3 (Cellsignaling, D13C9; 1:200), and anti-KRT8 (Origene, BP5075; 1:300). For secondary antibody incubation, the cells were incubated with Fluor 488- or 555- or 647-conjugated secondary antibodies 1 hour at room temperature. ProLong Diamond Antifade (DAPI included) was used to stain nuclei. The images were taken with a fluorescence confocal microscope (Leica). Paraffin sections were deparaffinized and then boiled in sodium citrate buffer (pH 6.0) for 20 min for antigen retrieval. Sections were blocked in 5% goat serum in TBST for 1 hour and incubated with primary antibodies at 4 °C overnight. The primary antibodies used in this experiment including anti-MMP2 (Cellsignaling, 40994; 1:200), anti-PL-1(Santa Cruz, sc-376436; 1:200) and anti-PTGS2 (Sigma, SAB2500267; 1:100-1:200). Then, the sections were incubated with fluorescence-conjugated secondary antibodies for one hour at room temperature. Nuclei were stained with DAPI (Invitrogen, D1306).

### Quantitative real-time PCR

Total RNA was extracted from cells using RNeasy Micro Kit (Qiagen) according to the manufacture’s protocol. First-strand cDNA was synthesized using the iScript cDNA Synthesis Kit (BIO-RAD). The qRT-PCR was performed using SYBR Green PCR Master Mix (BIO-RAD) with specific primers (Extended Data Tab. 2). Data were analyzed using the BIO-RAD software provided with the instrument. The relative gene expression values were calculated using the △△CT method and normalized to internal control GAPDH.

### IFNT activity analysis

IFNT activity was measured by an established IFN stimulatory response element-reporter assay [24]. Briefly, 5 - 10 × 10^5^ Madin-Darby bovine kidney cells (MDBK) that are stably transduced with an ISRE-Luc reporter were plated into a well of 96-well polystyrene plates with opaque walls and optically clear bottoms (Corning) and cultured in MDBK growth medium (high glucose DMEM, 10% FBS and 1% Pen/Strep) at 37°C for 4 hours. After removal of MDBK growth medium, 50 μL of sample or standard (Recombinant human IFN-α, IFNA: Millipore, IF007) were added. The standard curve was generated by a 1:3 serial dilution of IFNA. Cells were incubated at 37°C for 16 hours, then 50 μL One-Glow Luciferase reagent (Promega Corp; E6120) were added into each well, with a final volume of 100 μL. After mixture at a shaker platform for 10 minutes, the measurement was performed in a plate reader.

### TSCs lesion assay

bTSCs cells were grown to about 80% confluence in the bTSCs medium and dissociated with TrypLE. 5 × 10^6^ bovine TS cells were resuspended in 200 μL 1:1 of bTSC medium and Matrigel, and subcutaneously injected into 6-month-old non-obese diabetic (NOD)-severe combined immunodeficiency (SCID) mice. Lesions were collected at day 7 and 9, fixed in 4% PFA overnight at 4 °C for analysis.

### Karyotyping assay

bTSCs were incubated with bTSC medium containing 1 mL KaryoMAX colcemid solution (Gibco, 15212012) at 38.5 °C for 4-5 hours. Cells were then dissociated using 1 mL Trypsin (Gibco, 25200-056) at 38.5 °C and centrifuged at 300 × g for 5 min. The cells were resuspended in 1mL PBS solution and centrifuged at 400 × g for 2 min. The supernatant was aspirated and 500 μL 0.56% KCI was added to resuspend the cells. The cells were incubated for 15 min, then centrifuged at 400 × g for 2 min. 1 mL cold fresh Carnoy’s fixative (3:1 methanol: acetic acid) was added to resuspend the cells, followed by a 10 min incubation on ice. After centrifuge, 200 μL Carnoy’s fixative was added to resuspend the cells. Cells were dropped on the clean slides and air dried and soaked in a solution (1:25 of Giemsa stain (Sigma, GS500): deionized water) for 9 min. Slides were rinsed with deionized water and air dried. The images were taken by Leica DM6B at 1000× magnification under oil immersion.

### RNA sequencing analysis

Total RNA of bovine TSCs was extracted using RNeasy Micro Kit (Qiagen). Trophectoderm from day 7 blastocyst was isolated by placing embryos in a Petri dish with phosphate-buffered saline and performing microsurgery using a microblade under a microscope. The RNA-seq libraries were generated by using the Smart-seq2 v4 kit with minor modification from manufacturer’s instructions. Briefly, mRNA was captured and amplified with the Smart-seq2 v4 kit (Clontech). After AMPure XP beads purification, amplified RNAs were quality checked by using Agilent High Sensitivity D5000 kit (Agilent Technologies). High-quality amplified RNAs were subject to library preparation (Nextera XT DNA Library Preparation Kit; Illumina) and multiplexed by Nextera XT Indexes (Illumina). After purification of library with AMPure XP beads (Beckman Coulter), the concentration of sequencing libraries was determined by using Qubit dsDNA HS Assay Kit (Life Technologies). The size of sequencing libraries was determined by means of High Sensitivity D5000 Assay in at Tapestation 4200 system (Agilent). Pooled indexed libraries were then sequenced on the Illumina NovaSeq platform with 150-bp paired-end reads.

The Salmon tool [32] was applied to quantify the gene expression profile from the raw sequencing data, by using the Ensembl bovine genome annotation (ARS-UCD1.2). Transcript per million reads (TPM) was used as the unit of gene expression. The edgeR tool [33] was applied to identify differentially expressed genes. The TMM algorithm implemented in the edgeR package was used to perform normalization of the read counts and estimation of the effective library sizes. Differential expression analysis was performed by the likelihood ratio test implemented in the edgeR package. All the conventional statistical analyses were performed based on the R platform. The “cor.test” function was used to perform Spearman’s rank correlation test. Principal component analysis (PCA) on the gene expression profile was performed by using the “dudi.pca” function within the package “ade4”. All the heatmaps were plotted by the “heatmap.2” function within the package “gplots”. The gene ontology and pathway analysis were performed by means of the David tool [34].

In total, we sequenced two replicates of bTSCs, trophoblasts differentiated at day 2, 3, 4, 5 and 6, three replicates of whole blastocysts and day 7 trophectoderm cells selected from the same batch used for bTSCs derivation. The RNA-seq datasets of bovine day 14 trophectoderm [25], ESCs [27] and EPSCs [26] were downloaded from previous publications, respectively.

### ATAC-seq analysis

The ATAC-seq libraries from fresh cells were prepared as previously described [31]. Shortly, cells or embryos were lysed on ice, then incubated with the Tn5 transposase (TDE1, Illumina) and tagmentation buffer. Tagmentated DNA was purified using MinElute Reaction Cleanup Kit (Qiagen). The ATAC-seq libraries were amplified by Illumina TrueSeq primers and multiplexed by index primers. Finally, high quality indexed libraries were then pooled together and sequenced on Illumina NovaSeq platform with 150-bp paired-end reads.

The ATACseq analysis was followed our established analysis pipeline [31]. All quality assessed ATAC-seq reads were aligned to the bovine reference genome using Bowtie 2.3 with following options: –very-sensitive -X 2000 –no-mixed –no-discordant. Alignments resulted from PCR duplicates or locations in mitochondria were excluded. Only unique alignments within each sample were retained for subsequent analysis. ATAC-seq peaks were called separately for each sample by MACS2 with following options: –keep-dup all –nolambda –nomodel. The ATAC-seq bigwig files were generated using bamcoverage from deeptools. The ATAC-seq signals were visualised in the Integrative Genome Viewer genome browser. The annotations of genomic features, including transcription start sites, transcription end sites (TES), promoters, CDS, introns, 5′ UTR, 3′ UTR and intergenic regions were downloaded from UCSC genome browser. The enrichment of transcriptional factor motifs in peaks was evaluated using HOMER (http://homer.ucsd.edu/homer/motif/). For downstream analysis, we normalised the read counts by computing counts scaled by the number of sequenced fragments multiplied by one million (CPM).

### Whole genome bisulfite sequencing (WGBS) analysis

WGBS libraries were prepared using the TruSeq DNA Methylation Library Preparation Kit (Illumina). Briefly, genomic DNA was isolated using the DNeasy Blood & Tissue Kit (Qiagen) according to the manufacturer’s guide. Then, approximately 500 ng DNA were bisulfite treated using EZ DNA Methylation Kit (Zymo Research). Bisulfite-converted DNA was end-repaired, dA-tailed, and ligated with adapters following instructions of the TruSeq DNA Methylation Library Preparation Kit. Finally, high quality indexed libraries were then pooled and sequenced on Illumina NovaSeq platform with 150-bp paired-end reads.

WGBS data analysis was followed our established analysis pipelines [35, 36]. Briefly, WGBS raw data were removed first 12-bp at the 5’ end of both pairs, and reads with adapters and low-quality bases by using TrimGalore-0.4.3. The trimmed sequences were mapped to the bovine genome (ARS-UCD1.2) using Bismark. Uniquely mapped reads were then removed PCR duplicated reads and non-converted reads using deduplicate_bismark and filter_non_conversion. For avoiding the sequencing bias, only reads with 10x coverage was used in the downstream analysis. Methylation of each CpG site was calculated and methylation DNA methylation of each sample was calculated by averaging the consecutive genomic window of 300-bp tiles’ methylation. Differentially methylated regions (DMRs) were defined as common 300-bp tiles between two compared groups, which methylation levels ≥ 75% in one group, while ≤25% in another, and were significantly different by Fisher’s exact test (P-value ≤ 0.05, FDR ≤ 0.05). Hyper- and hypo-methylated tiles were those with DNA methylation levels ≥ 75% and ≤25%, respectively. The gene ontology and pathway analysis were performed by means of the David tool [34].

### Data availability

The raw FASTQ files and normalized read accounts per gene are available at Gene Expression Omnibus (GEO) (https://www.ncbi.nlm.nih.gov/geo/) under the accession number GSE2209252534.

## Author contributions

Y.W. and Z.J. conceptualized the idea and designed the research. Y.W. performed most of the experiments. L.Y. helped with TSC medium optimization and performed the TSC lesion assay. J.L., L.Z., H.M., Y.W., and H.S. performed genomic analysis. L. Z., H.M., and Y.W. performed embryo collection experiments. C.P.A. helped with TSC characterizations. Y.W., J.W., and Z.J. interpreted data and assembled the results. J.W. and Z.J. supervised the study. Y.W., J.W., and Z.J. wrote the manuscripts with inputs from all authors.

## Acknowledgements

This work was supported by the NIH Eunice Kennedy Shriver National Institute of Child Health and Human Development (R01HD102533) and USDA National Institute of Food and Agriculture (2019-67016-29863). J.W. is a New York Stem Cell Foundation–Robertson Investigator and Virginia Murchison Linthicum Scholar in Medical Research and funded by CPRIT (RR170076), NIH (GM138565-01A1 and OD028763), and Welch (854671).

## Extended Data Tables

**Extended Data Table 1.** Culture medias

|  |  |  | C1<br>[37] | C2 | C3<br>[38] | C4 | C5 | C6 | C7<br>[5] | C8<br>[13] | C9 | C10 | C11<br>[39] |
| --- | --- | --- | --- | --- | --- | --- | --- | --- | --- | --- | --- | --- | --- |
| Basal<br>mediums | N2B27 |  | ✓ |  | ✓ | ✓ | ✓ | ✓ |  | ✓ | ✓ | ✓ | ✓ |
|  | mTeSR |  |  | ✓ |  |  |  |  |  |  |  |  |  |
|  | RPMI1640 |  |  |  |  |  |  |  | ✓ |  |  |  |  |
| FBS or<br>KSR | FBS (%) |  |  |  |  |  |  |  | 20 |  |  |  |  |
|  | KSR (10%) |  |  |  |  |  |  |  |  |  | 10 | 10 |  |
| Factors | TGF-β1 | | ✓ | ✓ | | | | | | | | | |
|  | LIF |  | ✓ | ✓ | ✓ | ✓ | ✓ | ✓ |  | ✓ | ✓ | ✓ |  |
|  | bFGF |  | ✓ | ✓ |  |  | ✓ | ✓ |  |  | ✓ | ✓ | ✓ |
|  | FGF4 |  |  |  |  |  |  |  | ✓ |  |  |  |  |
|  | Heparin |  |  |  |  |  |  |  | ✓ |  |  |  |  |
| Small<br>molecule | WNT | CHIR99021 | ✓ | ✓ | ✓ | ✓ | ✓ | ✓ |  | ✓ | ✓ | ✓ | ✓ |
|  | MEK1/<br>MEK2 | PD0325901 | ✓ | ✓ |  |  |  |  |  |  | ✓ |  |  |
|  | JNK | SP600125 | ✓ | ✓ |  |  |  | ✓ |  |  |  |  |  |
|  | MAPK | SB203580 | ✓ | ✓ |  |  |  | ✓ |  |  |  |  |  |
|  | FGFR1 | SU5402 |  |  | ✓ | ✓ | ✓ |  |  |  |  |  |  |
|  | MEK | PD184352 |  |  | ✓ | ✓ | ✓ |  |  |  |  |  |  |
| | TGF-β | ActivinA | | | | ✓ | ✓ | | | | ✓ | ✓ | ✓ |
|  | muscarinic<br>M2/<br>histamine<br>H1 | DiM |  |  |  |  |  |  |  | ✓ |  |  |  |
|  | MMP | MiH |  |  |  |  |  |  |  | ✓ |  |  |  |
| Outgrowth rate (%) |  |  | 0<br>(0/3) | 0<br>(0/3) | 0<br>(0/3) | 0<br>(0/3) | 20<br>(1/5) | 0<br>(0/3) | 0<br>(0/3) | 89.3<br>(25/28) | 58.3<br>(7/12) | 66.7<br>(8/12) | 53.3<br>(8/15) |

C: combination; DiM: Dimethinedene maleate; MiH: Minocycline hydrochloride.

**Extended Data Table 2.**
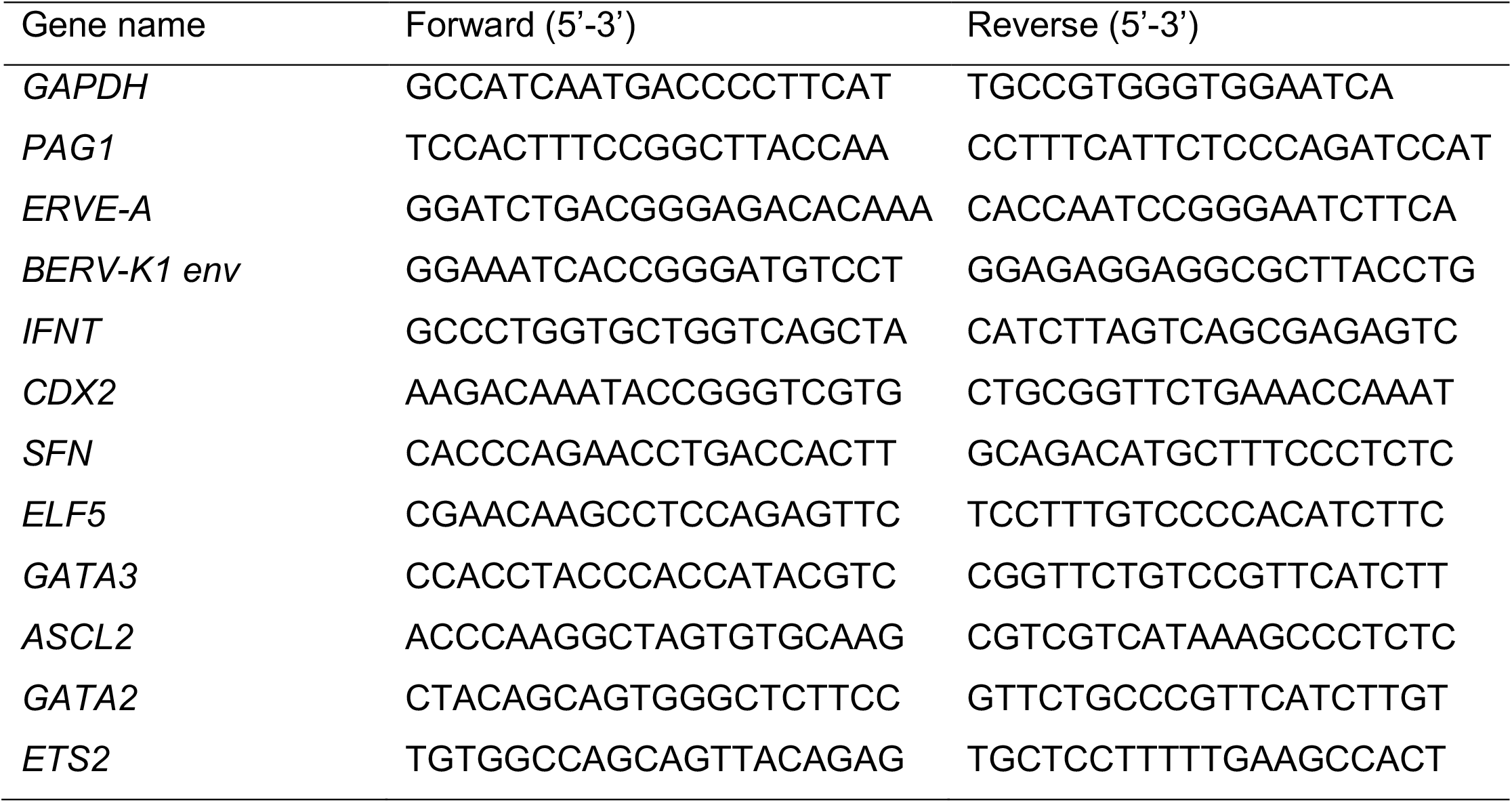
Primer lists

## Extended Data Figures

**Extended Data Fig 1.**
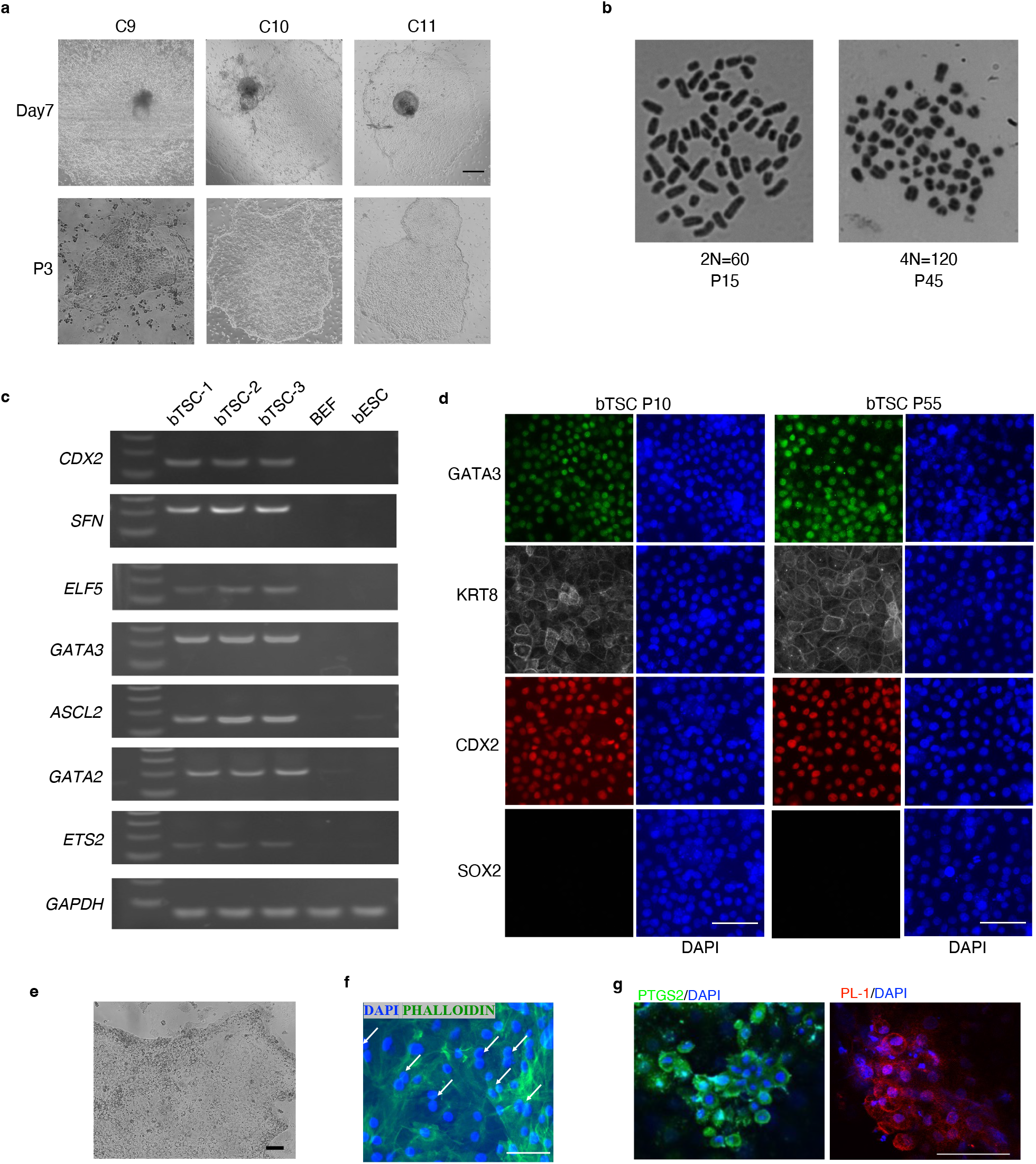
**a**, Representative images of the outgrowths of blastocysts after 7 days culture (top row) and cells after 3 passages (P3) (bottom row) in C9, C10 and C11 medium. Scale bar: 100μm. **b**, Karyotyping of bTSC at passage 15 and 45, respectively. **c**, RT-PCR analysis of bovine trophoblast marker genes (*CDX2, SFN, ELF5, GATA3, ASCL2, GATA2*, and *ETS2*) in bovine TSC. *GAPDH* serves as control. BEF: bovine embryonic fibroblast; bESC: bovine embryonic stem cells. **d**, Immunostaining for epiblast marker SOX2, and trophoblast marker (GATA3, KRT8, CDX2) in bTSC at passage 10 (P10) and passage 55 (P55) (Scale bar: 50μm). **e**, Bright filed image of differentiated-TSC. Scale bar: 50μm. **f**, Representative immunostaining images showing binucleation in differentiated-bTSC (P27). Scale bar: 100 μm. **g**, Representative immunostaining of mature trophoblast markers (PTGS2 and PL-1) in differentiated-bTSC (P27). Scale bar: 75 μm.

**Extended Data Fig 2.**
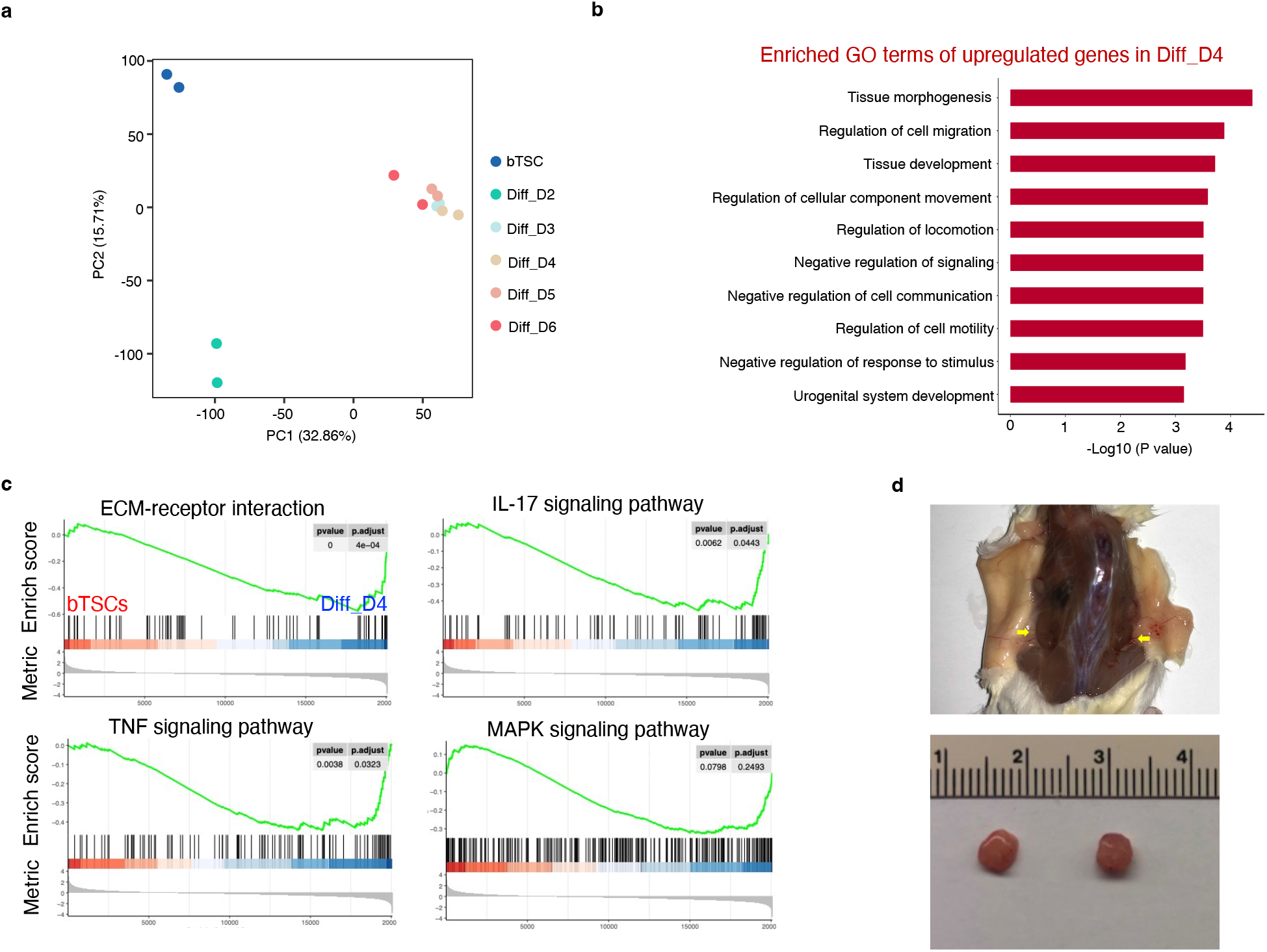
**a**, PCA analysis of transcriptomes of bTSC and differentiated-TSC at day 2, 3, 4, 5, and 6. **b**, Top 10 enriched gene ontology (GO) terms in Diff_D4 trophoblast compared with bTSC. **c**, Gene set enrichment analysis (GSEA) of bTSC and Diff_D4 trophoblast cells. Green line shows enrichment profile. Vertical black bars show where genes from a given gene set are located. **d**, NOD-SCID mice with tumor formed after bTSCs were injected (Top row). Tumors removed from mice after 9 days injection (Bottom row).

**Extended Data Fig 3.**
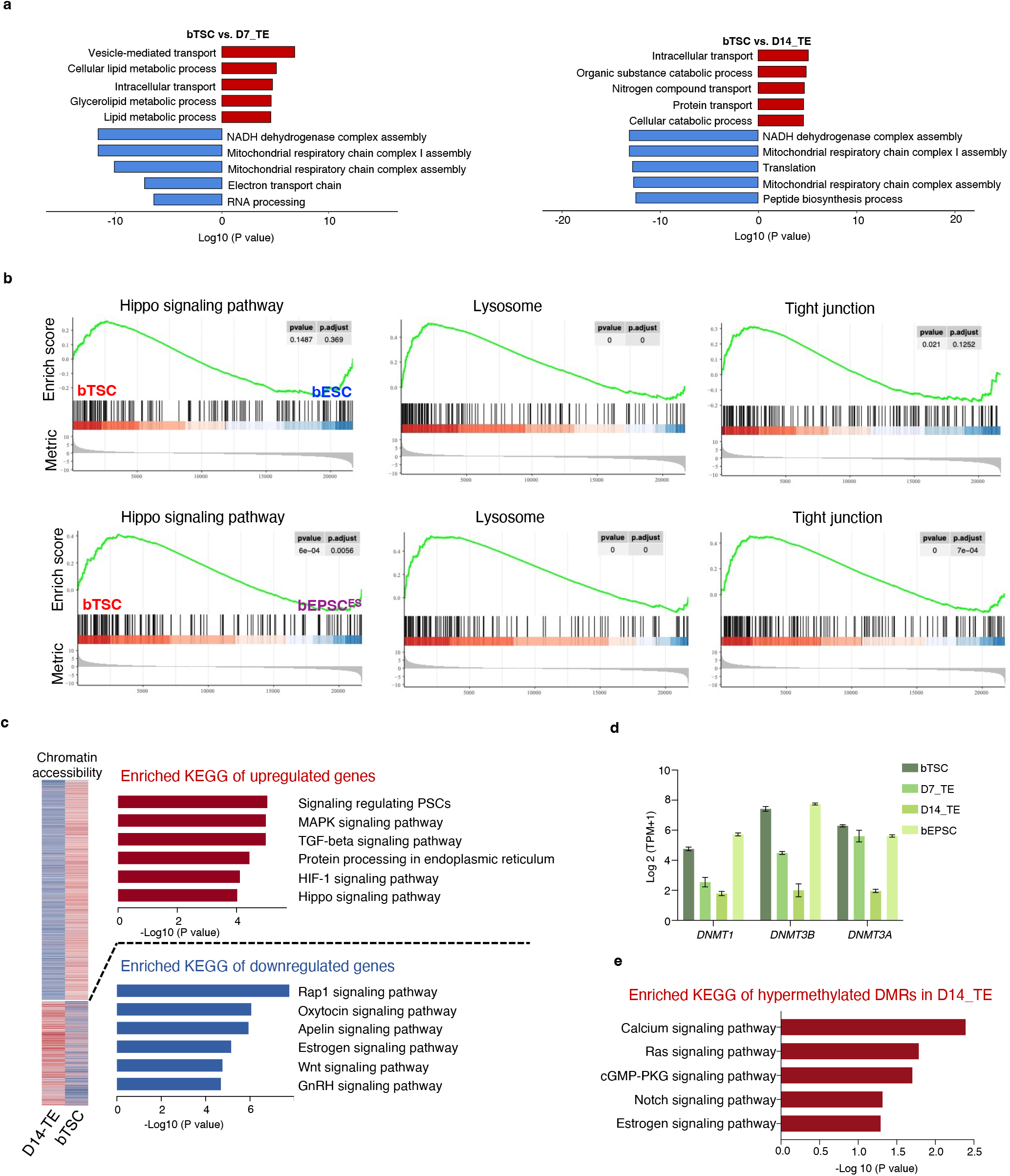
**a**, Top 5 enriched and depleted GO terms in bTSC compared to D7_TE or D14_TE. **b**, GSEA analysis of transcriptomes between bTSC and bESC and bEPSC^ES^. Genes with Hippo signaling pathway, lysosome and Tight junction were upregulated in bTSC. **c**, Pathways enriched in genes with more accessible or closed chromatin in bTSC compared to D14_TE. **d**, Expression levels of DNA methyltransferase (*DNMT1, DNMT3A*, and *DNMT3B*) in bTSC, D7_TE, D14_TE and bEPSC. **e**, Enriched pathways associated with genes annotated from hypomethylated DMRs in bTSC compared to D14_TE.

